# Chloride intracellular channel 4 (CLIC4) expression is transcriptionally regulated by crosstalk of the TGF-β, Wnt and Hedgehog signalling pathways

**DOI:** 10.1101/2021.06.29.449736

**Authors:** Christopher W. Wasson, Begoña Caballero-Ruiz, Jamel Mankouri, Gianluca Canettieri, Natalia A. Riobo-Del Galdo, Francesco Del Galdo

**Author notes:** Co-Corresponding Author: Francesco Del Galdo, Christopher W Wasson.

## Abstract

Chloride intracellular channel 4 (CLIC4) is a recently discovered driver of fibroblast activation in Scleroderma (SSc) and cancer-associated fibroblasts. CLIC4 expression and activity are regulated by TGF-β signalling through the SMAD3 transcription factor. In view of the aberrant activation of canonical Wnt and Hedgehog (Hh) signalling in fibrosis, we investigated their role in CLIC4 upregulation. Here, we show Wnt3a/β-catenin and Smoothened/GLI signalling cooperate with SMAD3 to regulate CLIC4 expression in normal dermal fibroblasts, and that inhibition of SMAD3 expression or activity abolishes Wnt and Hh-dependent CLIC4 induction. We further show that expression of the profibrotic marker α-smooth muscle actin strongly correlates with CLIC4 expression in dermal fibroblasts. Our data highlight novel mechanisms that regulate CLIC4 expression that present targetable pathways to prevent fibroblast activation in SSc and other fibrotic conditions.

## Introduction

Chloride intracellular channel 4 (CLIC4) is a small protein that can be found in soluble or membrane-associated forms and has multiple functions acting as a chloride channel and as a regulator of internal membrane and cytoskeletal dynamics and, therefore, plays an important role in a number of cellular functions including angiogenesis, proliferation, and inflammation. (1). CLIC4 translocates from the cytosol to the plasma membrane within seconds of serum stimulation or of activation of sphingosine-1-phosphate, muscarinic M3 and lysophosphatidic acid receptors (2, 3). One of its downstream functions is regulating G protein-coupled receptor activation. This leads to activation of RhoA and Rac1 as well as regulation of actin dynamics (3). CLIC4 has also been reported play an important role in cell division through acto-myosin contraction in the cleavage furrow during cytokinesis (4).

Fibroblast activation, characterised by expression of profibrotic genes such as α-smooth muscle actin (α-SMA) and collagens, is characteristic of scleroderma (SSc) and of cancer-associated fibroblasts (CAFs). CLIC4 is upregulated in both CAFs (5,6) and in SSc patient-derived fibroblasts (7). Upregulation of CLIC4 has been linked to increased fibroblast activation in response to TGF-β, the expression of pro-fibrotic genes (7), and the increased migration, invasion and epithelial to mesenchymal transition (EMT) of tumour cells associated with stromal CAFs (5). TGF-β induces CLIC4 nuclear translocation in a Schnurri-2 dependent manner (8). Nuclear translocation of CLIC4 is necessary for profibrotic gene expression and for prolonged TGF-β signalling through preventing the de-phosphorylation of SMADs 2 and 3 (8–9). CLIC4 is also a target gene of SMAD3 in fibroblasts, exerting a positive feedback loop to amplify TGF-β signalling (8).

It is now well established that during fibrosis there is a deregulation of Wnt and Hh signalling. Wnt signalling can be classified as canonical (β-catenin/TCF-dependent) or non-canonical (RhoA and calcium-dependent). Hh signalling has been reported to stimulate the Smoothened (SMO)/GLI axis in a canonical, primary cilium-dependent manner (10) and to trigger changes in RhoA/Rac1 activity and in voltage-dependent Kv channels by a non-canonical, GLI-independent pathway (11, 12). A wealth of studies in the literature indicates that Hh and Wnt signalling cascades crosstalk with the TGF-β pathway at multiple levels in positive feedback loops. Notably, SMAD3 is activated in gastric cancer cells stimulated with Shh (13) and the expression of GLI1 and GLI2 (the major Shh pathway transcription factors) is upregulated by TGF-β/SMADs (14). Further, TGF-β has been shown to stimulate canonical Wnt-3a/β-catenin signalling in a p38 MAPK-dependent manner in dermal fibroblasts (15). Concordantly, Henderson Jr. et al. have shown that the inhibition of β-catenin blocks α-SMA expression in response to TGF-β in alveolar epithelial cells (16). Direct interactions between SMAD3 and β-catenin have been identified at numerous promoter sites (17) including the GLI2 promoter (18). These findings are of interest in the context of SSc, where both Shh and Wnt-3a signalling are hyper activated and play important roles in the activation of SSc fibroblasts (19, 20).

Given the interconnection of TGF-β signalling with Hh and Wnt3a, we investigated the possibility that Hh and Wnt-3a signalling regulate CLIC4 expression in dermal fibroblasts. In this study, we show that stimulation of canonical Wnt3a and Hh signalling increases CLIC4 expression in dermal fibroblasts and this occurs in cooperation with TGF-β/SMAD3. In the context of SSc, we further demonstrate that CLIC4 expression can be supressed through inhibition of β-catenin and GLI1/GLI2-dependent transcription in patient dermal fibroblasts.

## Results

### Canonical Wnt and Hh signalling stimulate expression of CLIC4 in dermal fibroblasts

To investigate the role of morphogen signalling in the regulation of CLIC4 expression, normal human dermal fibroblasts were stimulated with TGF-β, Wnt-3a and SAG (a small molecule SMO agonist) for 48h. CLIC4 expression was investigated at both the protein and transcript level (Figure 1). Consistent with previous studies (7), TGF-β induced CLIC4 expression in dermal fibroblasts by 11-fold at mRNA (Figure 1A) and ~4-fold at protein level (Figure 1B). In the same experimental setting, Wnt-3a induced the expression of CLIC4 by 6-fold at mRNA and 3-fold at protein level. Similarly, SAG increased CLIC4 transcript by 7-fold at mRNA and 2.7-fold at protein level (Figure 1A, B). These data indicate that Wnt and Hh signalling activation may enhance CLIC4 expression similarly to what has been observed for TGF-β.

**Figure 1:**
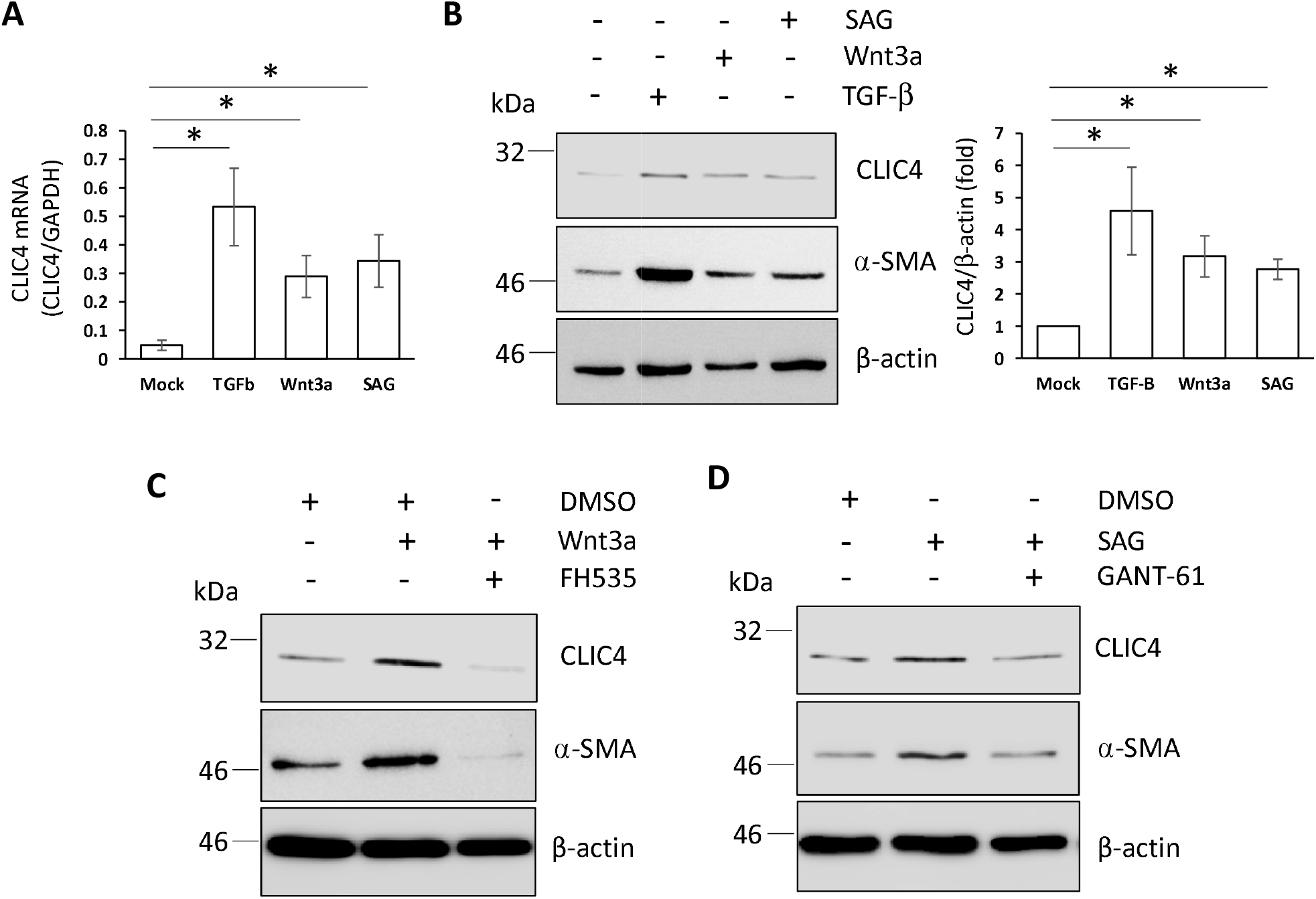
CLIC4 expression is regulated by a number of morphogens in dermal fibroblasts. Healthy dermal fibroblasts were grown in serum-depleted media and stimulated with TGF-β, Wnt3a and Smoothened agonist (SAG) for 48 h. RNA and protein were extracted from the cells. (A) CLIC4 transcript levels were analysed by q-RT-PCR. Graph represents the mean and standard error for 3 independent experiments. (B) CLIC4 and α-SMA protein levels were assessed by western blot. β-actin was used as a loading control. Densitometry analysis of CLIC4 blots. Graph represents the mean +/− standard error for 3 independent experiments. (C) Healthy dermal fibroblasts were grown in serum-depleted media and stimulated with Wnt3a for 48 h in the absence or presence of 10 uM FH535. CLIC4 and α-SMA protein levels were assessed by western blot. β-actin served as a loading control. (D) Healthy dermal fibroblasts were grown in serum-depleted media and stimulated with SAG for 48 h in the absence or presence of 10 uM GANT61. CLIC4 and α-SMA protein levels were assessed by western blot. β-actin served as a loading control. *p<0.05

We next sought to determine whether the observed effect was associated with activation of the canonical Wnt and Hh signalling pathways. To this end, we inhibited the transcriptional activity of β-catenin or the transcription factors GLI1 and GLI2, with the pharmacological β-catenin inhibitor FH535 and the GLI1/GLI2 inhibitor GANT61. Dermal fibroblasts treated with Wnt-3a in combination with FH535 suppressed the ability of Wnt-3a to induce CLIC4 upregulation at the protein level and reduced expression of CLIC4 below the basal level (Figure 1C). Similarly, GANT61 reversed the SAG-mediated increased in CLIC4 (Figure 1D). Strikingly, α-SMA expression closely correlated with CLIC4 expression levels in all conditions, confirming a previous report that CLIC4 silencing abolished α-SMA expression in fibroblasts (5). These data suggest that, in addition to SMAD2-3, β-catenin and the GLI transcription factors, contribute to CLIC4 expression in dermal fibroblasts.

### β-catenin drives CLIC4 overexpression in SSc patient fibroblasts

We have previously reported that CLIC4 is overexpressed in SSc patient fibroblasts through the activity of the SMAD3 transcription factors (7). Since Wnt/β-Catenin signalling is dysregulated in SSc fibroblasts (20) we set out to determine if Wnt/β-Catenin contribute to the high expression of CLIC4 in SSc fibroblasts (7). To investigate a role for Wnt/β-Catenin signalling, healthy and SSc patient-derived dermal fibroblasts were treated with the β-Catenin inhibitor FH535, and CLIC4 expression was assessed. FH535 treatment led to a partial reduction in CLIC4 levels at both the transcript and protein levels in SSc fibroblasts, but had no effect on CLIC4 expression in healthy dermal fibroblasts (Figure 2A). This follows a similar trend to α-SMA levels (Figure 2A-B) further confirming a previous report that CLIC4 silencing abolished a-SMA expression in fibroblasts. These data suggest that enhanced β-catenin signalling contributes to regulate CLIC4 expression in SSc.

**Figure 2:**
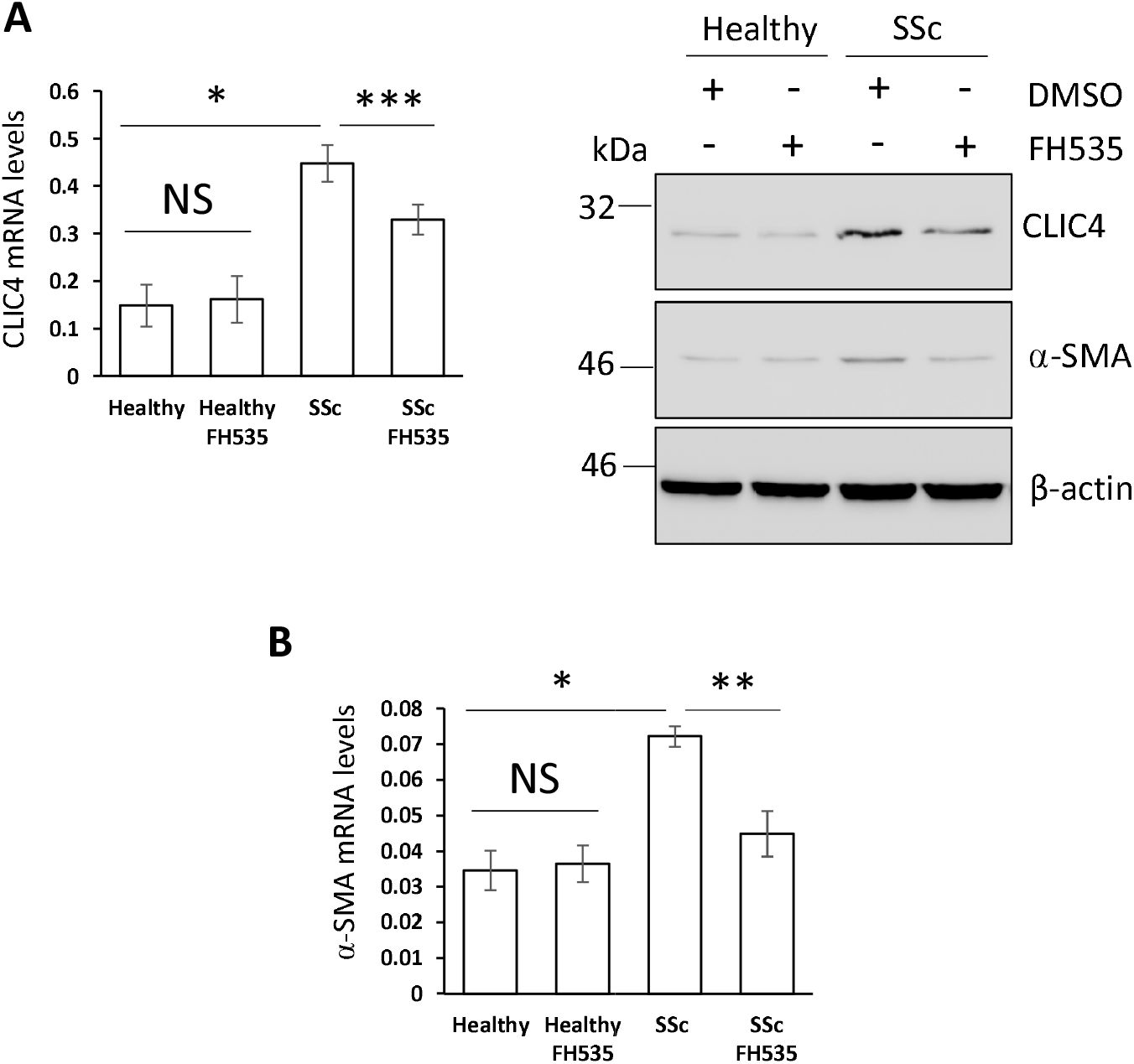
β-Catenin contributes to the enhanced CLIC4 expression in SSc fibroblasts. RNA and protein were extracted from healthy and SSc dermal fibroblasts treated with 10 uM FH535 or DMSO (vehicle) for 48 h. (A) CLIC4 transcript levels were assessed by q-RT-PCR. Graph represent the mean +/− standard error for 3 independent experiments. CLIC4 and α-SMA protein levels were assessed by western blot. β-actin served as a loading control. (B) α-SMA transcript levels were assessed by q-RT-PCR. Graph represent the mean +/− standard error for 3 independent experiments. *p<0.05, **p<0.01, ***p<0.001

### Enhanced CLIC4 levels in SSc fibroblasts are mediated by GLI2 expression

Dysregulated Hh signalling is a characteristic feature of SSc fibroblasts (19). In view of our in vitro data (Figure 1) we hypothesis that GLI transcription factors may contribute to the increased CLIC4 levels observed in SSc fibroblasts. To test this hypothesis, healthy and SSc patient-derived dermal fibroblasts were treated with the GLI1 and GLI2 inhibitor GANT61, followed by assessment of CLIC4 expression. Similarly, to what we observed for Wnt/β-Catenin pathway, GANT61 treatment led to a reduction in CLIC4 transcript and protein levels in SSc fibroblasts, but had no effect on CLIC4 expression in healthy fibroblasts (Figure 3A-B). These data suggest a role for GLI transcription factors in the overexpression of CLIC4 in SSc fibroblasts.

**Figure 3:**
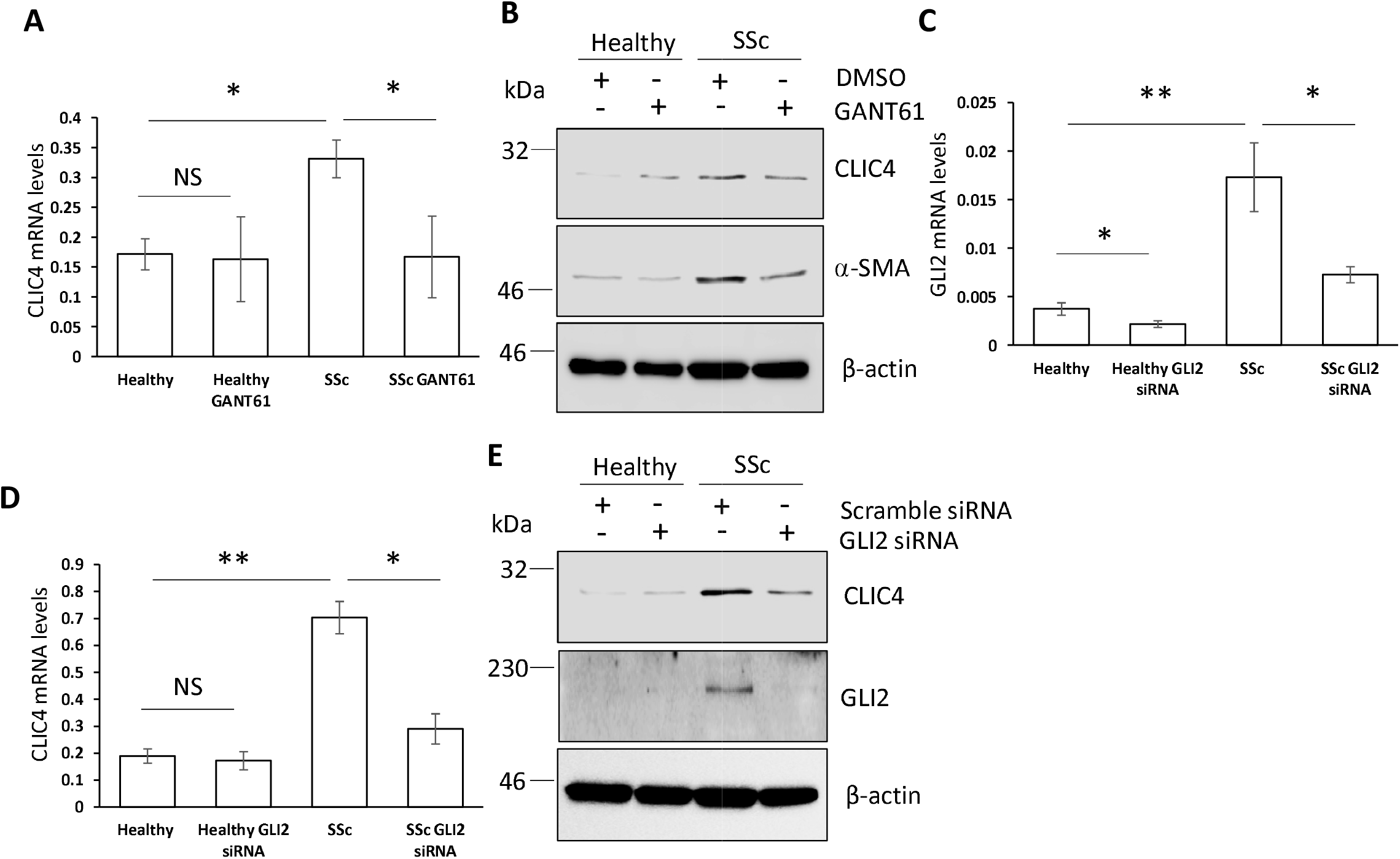
GLI2 is an important mediator of enhanced CLIC4 levels in SSc fibroblasts. (A) CLIC4 transcript levels from healthy and SSc fibroblasts treated or not with 10 uM GANT61 for 24 h were assessed by q-RT-PCR. Graph represents the mean +/− standard error for 3 independent experiments. (B) CLIC4 and α-SMA protein levels were assessed by western blot. β-actin served as a loading control. Healthy and SSc fibroblasts were transfected with siRNA specific to GLI2 or scramble control siRNA. (C) GLI2 transcript levels were assessed by q-RT-PCR. Graph represents the mean +/− standard error of 3 independent experiments. (D) CLIC4 mRNA levels were quantified in the same samples from (C). (E) CLIC4 and GLI2 protein levels from healthy and SSc fibroblasts treated with GLI2 or scrambled siRNA were assessed by western blot. β-actin served as a loading control. *p<0.05, **p<0.01

GLI2 is the main mediator of the canonical Hh pathway in response to SHh or to a direct SMO agonist, while GLI1 is induced in response to GLI2 and GLI3 activation (10). Since the SMO agonist SAG enhanced CLIC4 expression (Figure 1), and GANT61 inhibits both GLI1 and GLI2, we investigated the specific contribution of GLI2 using siRNA silencing approaches in healthy and SSc fibroblasts. GLI2 silencing was confirmed by qPCR and western blot analysis (Figure 3C & E) and this was found to significantly reduce the expression of CLIC4 in SSc fibroblasts, but not in healthy cells (Figure 3D & E). This data confirmed a key role for GLI2 in the overexpression of CLIC4 observed in SSc fibroblasts.

### Wnt3a and Hh signalling cooperate with SMAD3 to enhance CLIC4 expression

The evidence above shows that the canonical Wnt and Hh signalling pathways contribute to the elevated levels of CLIC4 observed in SSc fibroblasts, in addition to the previously described role for the TGF-β signalling pathway (7). These three pathways may regulate CLIC4 independently or in cooperation. Previous studies have shown both the Wnt and Hh signalling pathways interact with elements of the TGF-β signalling pathway in a number of cell types (13–18). In order to establish the hierarchical relationship among TGF-β and Wnt signalling in dermal fibroblasts, we investigated the effect of SMAD3 inhibition with siRNA or a small molecule inhibitor on Wnt/β-catenin-dependent transcription, as determined using a TOP Flash-luciferase reporter. As seen in Figure 4A, reduction in SMAD3 protein level to ~ 50% in healthy dermal fibroblasts led to a ~50% inhibition of TOP Flash luciferase activity in response to Wnt3a, while a scrambled siRNA duplex did not have any effect. In support, the SMAD3 inhibitor SIS3 reduced Wnt3a-mediated stimulation of TOP Flash-luciferase activity in a dose dependent manner in healthy dermal fibroblasts (Figure 4B). Therefore, high basal TGF-β/SMAD3 signalling in healthy dermal fibroblasts may enhance Wnt3a/β-catenin signalling.

**Figure 4:**
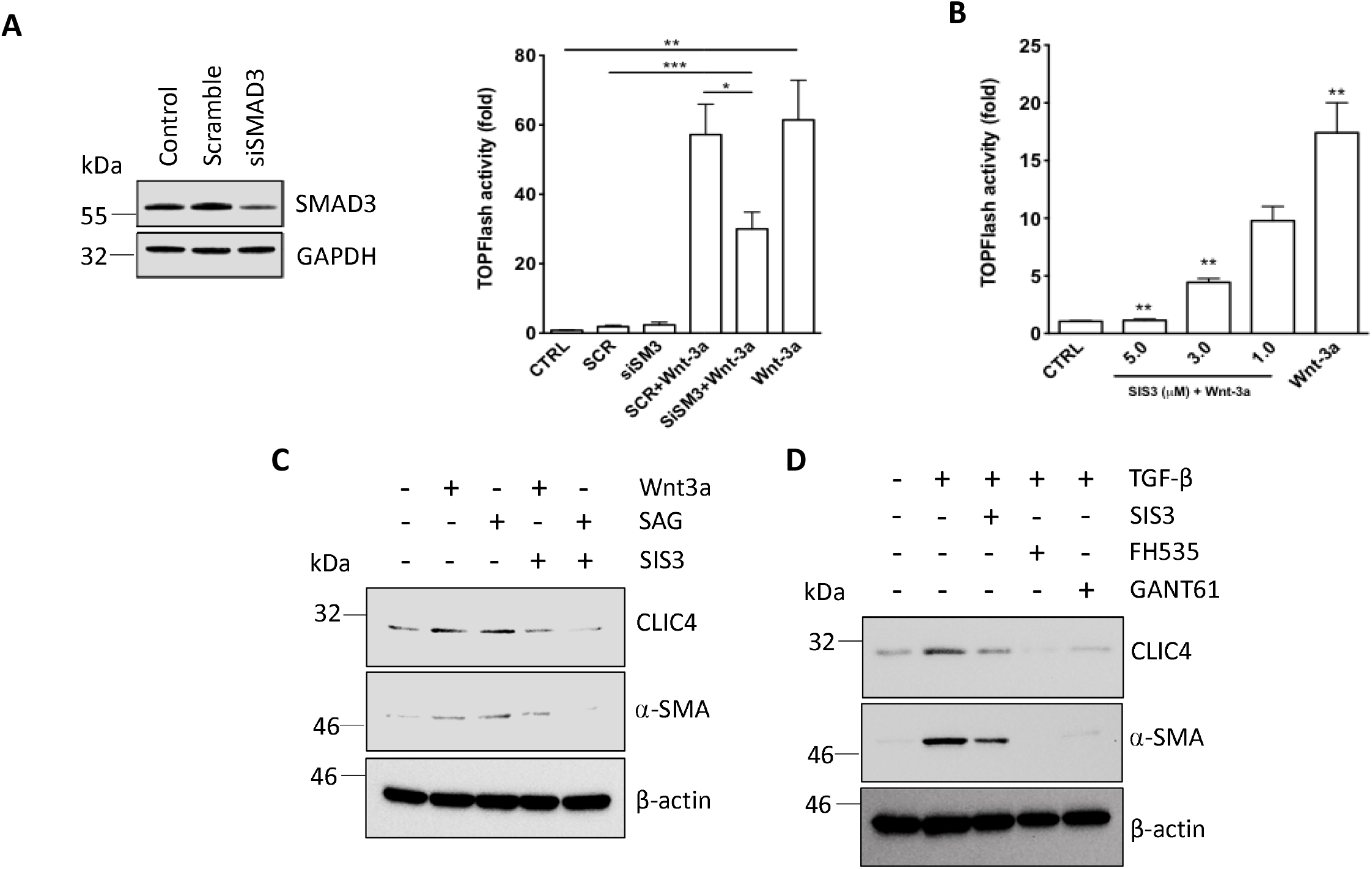
SMAD3 is required for CLIC4 expression induced by Wnt3a and a SMO agonist. Healthy dermal fibroblasts were transfected with siRNA specific to SMAD3 (siSMAD3) or scramble control siRNA (SCR) for 48 h, followed by transfection with the TOPFlash-Firefly and CMV-Renilla luciferase reporters for additional 24 h prior to treatment with Wnt3a (100ng/ml) for an additional 24 hours. (A) Left: SMAD3 protein levels were assessed by western blot. GAPDH was used as a loading control. Right: Relative TOPFlash reporter activity was determined by the dual luciferase assay. Data shown as mean ± SEM, n=4; Mann-Whitney test for unpaired samples. (B) Healthy dermal fibroblasts were treated with SIS3 (0-10 μM) for 1 h prior to Wnt3a (100 ng/ml) stimulation for 24 h. Relative TOPFlash reporter activity was determined by the dual luciferase assay. Data shown as mean ± SEM, n=4; Mann-Whitney test for unpaired samples. (C) CLIC4 and α-SMA protein levels in healthy fibroblasts stimulated with 100 ng/ml Wnt3a or 100 nM SAG for 48 h in the absence of presence of the DIM SIS3. β-actin served as a loading control. (D) CLIC4 and α-SMA protein levels in healthy fibroblasts stimulated with 10 ng/ml TGF-b for 48 h in the absence of presence of the 1□M SIS3, 10 uM FH535, or 10 uM GANT61. β-actin served as a loading control. *P<0.05, **P<0.01, ***P<0.001

To investigate if CLIC4 upregulation in response to Wnt3a/Hh pathway activation is also potentiated by SMAD3, we treated healthy dermal fibroblasts with the Wnt3a and SMO agonist SAG in combination with SIS3. Interestingly, SIS3 blocked both Wnt3a and SAG-induced CLIC4 and α-SMA expression (Figure 4C). Finally, we investigated whether β-catenin and GLI2 were mediators of TGF-β ability to induce CLIC4 expression. Healthy dermal fibroblasts were stimulated with TGF-β in combination with the SMAD3 inhibitor SIS3, the β-catenin inhibitor FH535, or the GLI1/2 inhibitor GANT61. Remarkably, blocking β-catenin or GLI1/2 inhibited TGF-β mediated CLIC4 and α-SMA expression to a greater extent than the SMAD3 inhibitor (Figure 4D). These results suggest that high basal SMAD3 activation in SSc fibroblasts cooperates with Wnt and Hh signalling to stimulate α-SMA expression via upregulation of CLIC4.

### Regulation of CLIC4 expression by SMADs and GLI signalling crosstalk is not limited to fibroblasts

In a parallel approach, we investigated if the observed regulation of CLIC4 by crosstalk of TGF-β and Hh signalling is specific to dermal fibroblasts or it is a widespread phenomenon. To this end, we used SW620 cells, a colon cancer cell line without mutations in the Hh pathway in which we engineered a pathogenic PTCH1 mutation using CRISPR/Cas9 technology. The phenotypic behaviour and whole transcriptome changes resulting from this mutation are described elsewhere (manuscript in preparation). Of interest in this study, transcriptome analysis of TGF-β and Hh signalling components in control (wt PTCH1) cells transfected with a scrambled gRNA and mutant PTCH1 (PTCH1^mut^) cells by RNA-seq revealed that PTCH1^mut^ cells expressed higher RNA levels of SMAD2, SMAD3 and SMAD4, higher levels of SMO, lower levels of GLI3, and higher levels of CLIC4 (Table 1). Higher levels of SMAD2/3/4 suggest an upregulation of SMAD-dependent TGF-β signalling, while upregulation of SMO together with a significant reduction in GLI3 expression, which acts mainly as a repressor of GLI1 and GLI2 target genes, suggest increased GLI transcriptional activity. In agreement with the RNA-seq data, PTCH1^mut^ cells had significant upregulation of GLI1 (4-fold; p<0.01) and CLIC4 (30-fold, p<0.01) transcript compared to control cells, which were also evident at protein level (Figure 5A-C). Treatment of PTCH1^mut^ cells with GANT61 simultaneously reduced both GLI1 expression and upregulation of CLIC4 by ~50% and ~75% respectively (Figure 5B). Western blot analysis of protein lysates confirmed the mRNA data. In the same experimental settings, we analysed the effect of the TGF-β receptor inhibitor SD208. Similar to GANT61 effects, SD208 suppressed the increased expression of CLIC4 observed in PTCH1^mut^ by 70% at mRNA level, with slightly less apparent effect at protein level (Figure 5C). Taken together this data supports a widespread crosstalk of Hh and TGF-β signalling in different cell types, that plays an essential role in regulating CLIC4 expression (Figure 5D).

**Table 1.**
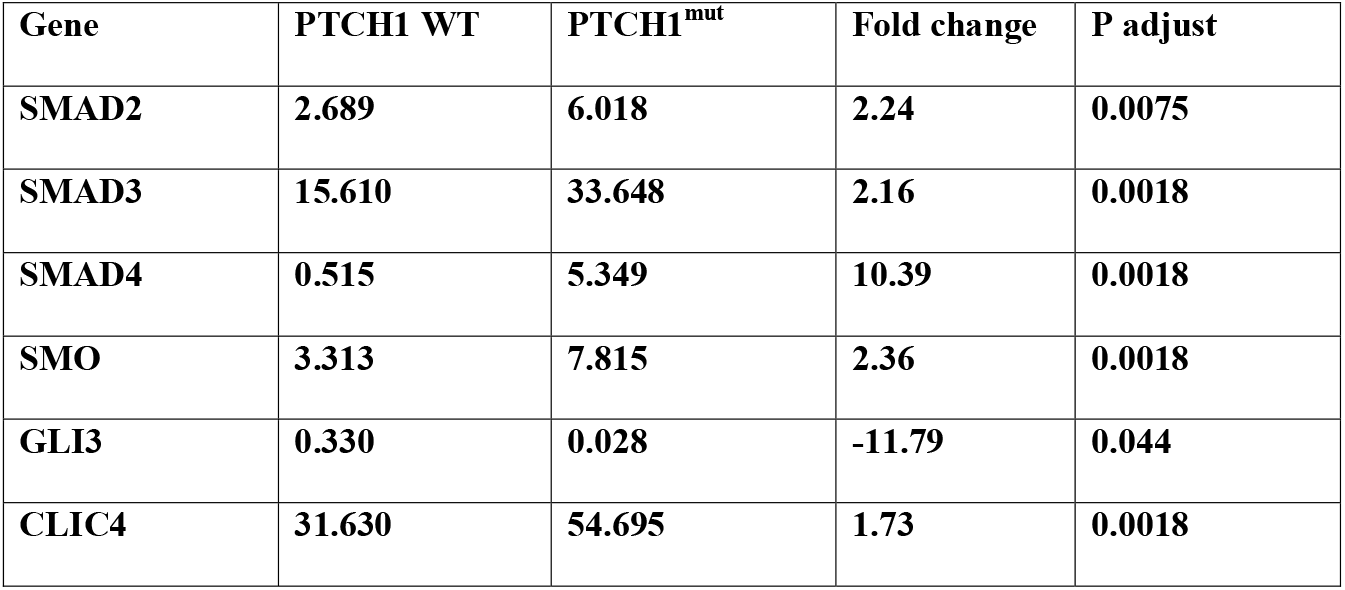
mRNA expression (RPKM values) of TGF-β and Hh pathway mediators in isogenic PTCH1 WT and PTCH1^mut^ SW620 cells.

| **Gene** | **PTCH1 WT** | **PTCH1<sup>mut</sup>** | **Fold change** | **P adjust** |
| --- | --- | --- | --- | --- |
| **SMAD2** | **2.689** | **6.018** | **2.24** | **0.0075** |
| **SMAD3** | **15.610** | **33.648** | **2.16** | **0.0018** |
| **SMAD4** | **0.515** | **5.349** | **10.39** | **0.0018** |
| **SMO** | **3.313** | **7.815** | **2.36** | **0.0018** |
| **GLI3** | **0.330** | **0.028** | **-11.79** | **0.044** |
| **CLIC4** | **31.630** | **54.695** | **1.73** | **0.0018** |

**Figure 5:**
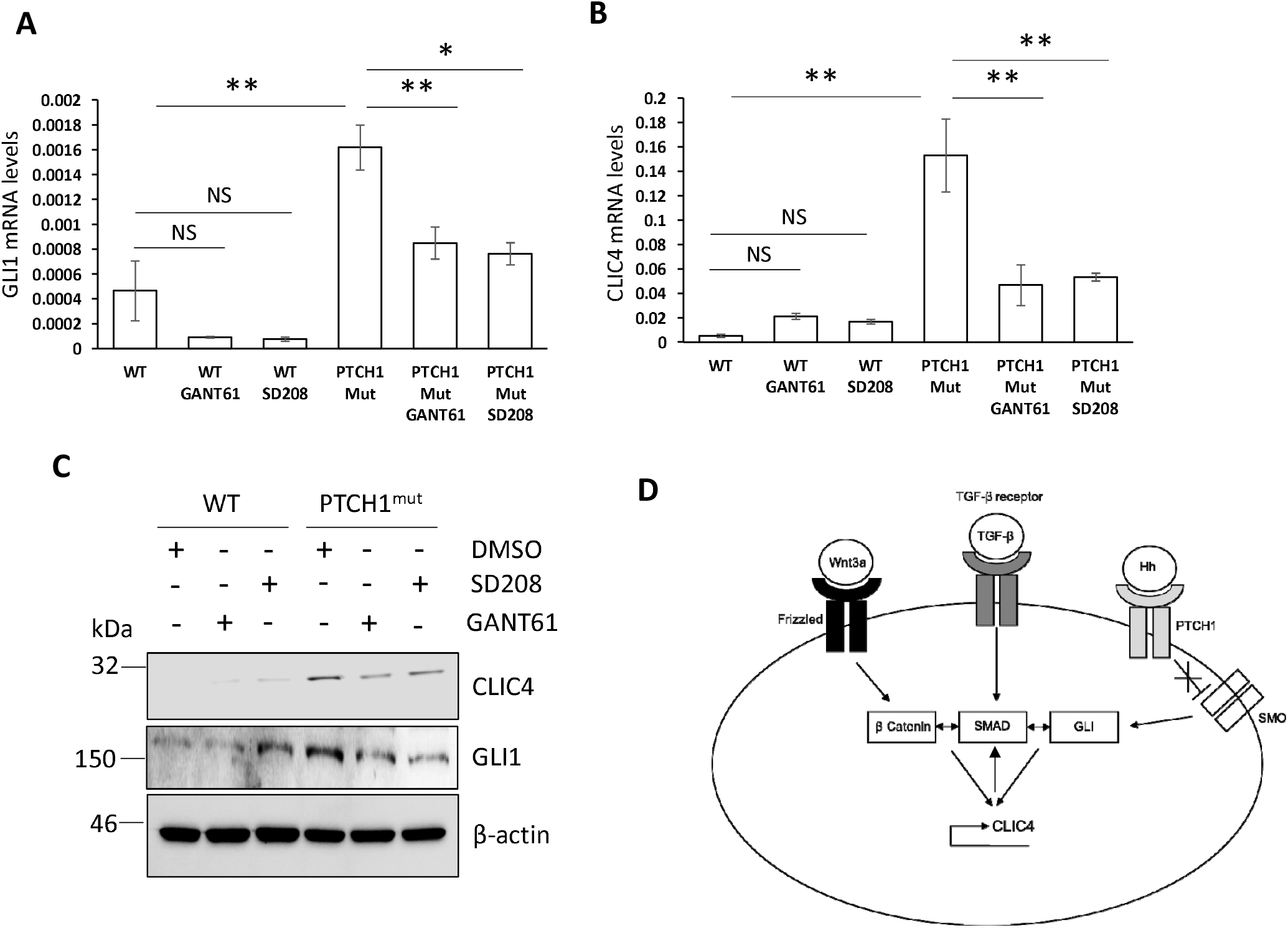
Regulation of CLIC4 expression by TGF-β and Hh signaling is conserved in other cell types. Isogenic SW620 cells expressing endogenous WT PTCH1 or mutated PTCH1 (PTCH1^mut^) were treated with 20 uM GANT61, 10 uM SD208, or DMSO for 24 h. RNA and protein were extracted from the cells. GLI1 (A) and CLIC4 (B) transcript levels were assessed by qPCR after 24 h in the presence of the inhibitors. Graphs represent the mean +/− standard error of 3 independent experiments. (C) CLIC4 and GLI1 protein levels were analysed by western blot. β-actin was used as a loading control. (D) Schematic of TGF-β, Wnt3a and Hh mediated regulation of CLIC4 expression. *P<0.05, **P<0.01

## Discussion

*CLIC4* is a target gene of TFG-β signalling and a key mediator of SMAD2/3-mediated transcription by preventing their dephosphorylation in the nucleus (5–9). CLIC4 expression is upregulated by TGF-β in dermal fibroblasts of SSc patients (7). In our previous study, we demonstrated that full expression of the myofibroblast phenotype – expression of α-SMA and collagens – requires CLIC4 activity as a chloride channel. In this study, we show that, in addition to its regulation by TGF-β, CLIC4 expression requires simultaneous activation of canonical Wnt and Hh signalling, acting through their cognate transcription factors β-catenin and GLI1/GLI2, respectively. Aberrant activation of these pathways in SSc dermal fibroblasts, in part as a result of excessive TGF-β/SMAD signalling, appears to be necessary for CLIC4 upregulation and consequent α-SMA expression.

The mechanistic basis of the functional cooperation among SMAD3, β-catenin and GLI1/GLI2 is not fully understood. SMAD3 has been shown to induce nuclear translocation of β-catenin in mesenchymal stem cells (21) and upregulates some Wnt family members in vascular smooth muscle cells (22), which is expected to stimulate β-catenin function as a transcriptional coactivator of TCF. SMAD2/3 and β-catenin/TCF4 have been reported to cooperate to stimulate GLI2 transcription by binding to specific elements in the *GLI2* promoter (18). Gli2 upregulation leads to a small but significant build-up of GLI1, a strong transcriptional activator. Thus, a possible explanation of our data is that tonic TGF-β in SSc fibroblasts increases Wnt/β-catenin signalling, and that both SMAD3 and β-catenin converge at the GLI2 promoter to induce upregulation of GLI2 and GLI1, this in turn may drive CLIC4 transcriptional upregulation. This model is supported by the almost complete repression of CLIC4 expression by the GLI1/2 inhibitor GANT61 and by GLI2 siRNA in SSc fibroblasts. This model also explains the findings in another cell type, in which dysregulation of Hh and TGF-β signalling by a mutation in PTCH1 leads to CLIC4 induction. SMAD3 and β-catenin have also been known to interact at a number of additional promoters (17); therefore, it is also possible they could directly interact at the CLIC4 promoter in parallel to a requirement for GLI1 or GLI2.

The use of pharmacological inhibitors is subjected to potential non-specific effects or cytotoxicity. However, GANT61 and GLI2 siRNA exerted a comparable inhibition of CLIC4 expression. The less profound effect of the SMAD3 inhibitor SIS3 on CLIC4 induction by TGF-β could be explained by redundant function of SMAD2, which is not targeted by the isoform-specific SMAD inhibitor. Therefore, our findings strongly support a sequence of signalling events that cooperate to drive CLIC4 expression.

Some findings suggest that the crosstalk between Hh and TGF-β signalling can be bidirectional. In PTCH1^mut^ cells, dysregulated Hh signalling is associated with upregulation of SMAD2, 3, and 4 mRNA, as was also shown for SMAD3 in gastric cancers (13). This could in turn act as a positive feedback loop to further increase and/or maintain CLIC4 levels.

TGF-β, Wnt/β-catenin and Hh are dysregulated in SSc fibroblasts and play important roles in SSc fibroblast activation. In this study, we show that a converging hub of all three morphogen pathways is the expression of CLIC4, which is a necessary mediator of several actions of TGF-β induced cellular events, including expression of profibrotic markers.

β-Catenin and the GLI transcription factors may also drive CLIC4 expression in cancer-associated fibroblasts (CAFs). Previous studies have shown both Hh and Wnt/β-catenin signalling contribute to the phenotypes of CAFs. For example, pancreatic derived CAFs show elevated levels of SMO and GLI1 expression, which was abolished by SMO silencing (23), whilst melanoma associated CAFs exhibit reduced ECM production and paracrine signalling when β-catenin is inhibited (24). Therefore, in addition to the well characterised role of TGF-β, Wnt-3a and Hh pathways may further drive CLIC4 expression in CAFs as in SSc fibroblasts and colon cancer cells shown here.

Overall, the convergence of TGF-β, Wnt/β-catenin and SHH signalling pathways on CLIC4 supports the role of CLIC4 as therapeutic target in loss of homeostasis during fibroais or desmoplastic reaction to cancer.

## Materials and Methods

### Patient cell lines

Full thickness skin biopsies were surgically obtained from the forearms of four adult patients with recent onset SSc, defined as a disease duration of less than 18 months from the appearance of clinically detectable skin induration. All patients satisfied the 2013 ACR/EULAR criteria for the classification of SSc and had diffuse cutaneous clinical subset as defined by LeRoy et al (25). All participants provided written informed consent to participate in the study. Informed consent procedures were approved by NRES-011NE to FDG. Fibroblasts were isolated and established as previously described (20, 26). Primary cells were immortalized using human telomerase reverse transcriptase (hTERT) to produce healthy control hTERT and SSc hTERT (20).

### Cell culture

hTERT patient fibroblasts were maintained in Dulbecco’s modified Eagle medium (DMEM) (Gibco) supplemented with 10% FBS (Sigma) and penicillin-streptomycin (Sigma). For indicated treatments fibroblasts were treated with vehicle (DMSO, Sigma) or equal volumes of GANT61 (10μM, Selleckchem), SIS3 (1-5μM, Selleckchem) and FH535 (10μM, Tocris) for 48 h in a humidified incubator at 37C and 5% CO_2_.

SW620 cells were obtained from ATCC. Cells were engineering by CRISPR/Cas9 to introduce a truncation in the *PTCH1* coding sequence (PTCH1^mut^) using short guide RNA (sgRNA) targeting the region of interest (D1222fs) or an irrelevant scrambled sequence (WT PTCH1) and selected using puromycin for 72 h, followed by generation of single cell-derived clones by the limiting dilution method. Individual colonies were sequence-verified and expanded in DMEM supplemented with 10% FBS and penicillin-streptomycin in a humidified incubator at 37C and 5% CO_2_. For the indicated treatments, SW620 clones were incubated with GANT61 (20μM, Stratech Scientific Ltd) or SD208 (10μM, Selleckchem) or equal volumes of DMSO for 24 h, followed by lysis and extraction of total RNA for further processing or by direct lysis in RIPA buffer (Thermo Scientific) containing protease and phosphatase inhibitor cocktails for western blotting (Thermo Scientific)(SantaCruz).

### Morphogen pathway stimulation

Healthy dermal fibroblasts were serum starved for 24 h in DMEM containing 0.5% FBS and stimulated with 10 ng/ml TGF-β (R&D systems), 100 ng/ml Wnt-3a (R&D systems) and 100 nM SMO agonist (SAG) for 24-48 h.

### TOP Flash TCF/LEF–firefly luciferase reporter

Fibroblasts were transfected with the TOP Flash TCF/LEF–firefly luciferase reporter vector for 24 hours (gift from Prof. Randall. Moon, University of Washington, Seattle, WA). pCMV-Renilla luciferase vector was co-transfected as a transfection control. These fibroblasts were then grown in serum depleted media for 24 h prior to stimulation. Cells were then harvested and luciferase activity was measured using the dual luciferase reporter assay reagents (Promega), as per the standard protocol, and detected using a luminometer (Berthold Mithras). To account for transfection efficiency, the Firefly luciferase activity was normalised to Renilla luciferase activity and expressed as the percentage change compared to control.

### siRNA transfections

A pool of four siRNAs specific for different regions of GLI2, SMAD3 or a negative control scrambled siRNA (Qiagen) were transfected into fibroblasts using Lipofectamine 2000 (Thermo Fisher). Fibroblasts were incubated for 48-72 h prior to harvesting.

### Western blotting

Total proteins were extracted from fibroblasts in ice-cold RIPA buffer and resolved by SDS-PAGE (10-15% Tris-Glycine). Proteins were transferred onto Hybond nitrocellulose membranes (Amersham biosciences) and probed with antibodies specific for alpha smooth muscle actin (Abcam), CLIC4 (Santa Cruz), GLI1 (Santa Cruz), GLI2 (R&D systems), SMAD3 (Cell signalling) and β-Actin (Sigma). Immunoblots were visualized with species-specific HRP conjugated secondary antibodies (Sigma) and ECL (Thermo/Pierce) on a Biorad ChemiDoc imaging system.

### Quantitative Real time PCR

RNA was extracted from cells using commercial RNA extraction kits (Zymo Research). RNA (1ug) was reverse transcribed using cDNA synthesis kits (Thermo). QRT-PCRs were performed using SyBr Green PCR kits on a Thermocycler with primers specific for alpha SMA (Forward: TGTATGTGGCTATCCAGGCG; Reverse: AGAGTCCAGCACGATGCCAG), CLIC4 (Forward: CATCCGTTTTGACTTCAGTGTTG; Reverse: AGGAGTTGTATTTAGTGTGACGA), GLI1 (Forward: GGACCTGCAGACGGTTATCC; Reverse: AGCCTCCTGGAGATGTGCAT), GLI2 (Forward: TTTATGGGCATCCTCTCTGG; Reverse: TTTTGCATTCCTTCCTGTCC) and GAPDH (Forward: ACCCACTCCTCCACCTTTGA; Reverse; CTGTTGCTGTAGCCAAATTCGT). Data were analysed using the ΔΔCt method using GAPDH as a housekeeping control gene.

### Statistical analysis

Data are presented as the mean ± standard error. Statistical analysis was performed using a two-tailed, paired Student’s t-test.

## Data Availability

All data generated or analysed during this study are included in the published article. Datasets are available from the corresponding author upon reasonable request.

## Acknowledgement

**C.W.W** is supported by Susan Cheney Scleroderma fellowship. **B.C.R. is supported by an international doctoral fellowship of the Sapienza University of Rome. N.R.DG**. is supported by the Biotechnology and Biological Sciences Research Council (BBSRC) project grant BB/S01716X/1. **F.DG** is supported by the National Institute for Health Research (NIHR) Leeds Biomedical Research Centre (BRC). The views expressed are those of the author and not necessarily those of the NIHR or the Department of Health and Social Care.

## Author Contributions Statement

**C.W.W:** Conceptualization, Investigation, Writing: Original draft preparation, Writing: Reviewing and editing. **B.C.R:** Investigation, Writing: Reviewing and editing. **J.M:** Resources, Investigation, Writing: Reviewing and editing. **G.C:** Resources, Investigation. **N.R.DG:** Resources, Investigation, Writing: Reviewing and editing. **F.DG:** Conceptualization, Funding Acquisition, Writing: Reviewing and editing

## References

1. Rao SG, Ponnalagu D, Patel NJ, Singh H. Three decades of chloride intracellular channel proteins: from organelle to organ physiology. Curr Protoc Pharmacol 2018; 80:11.21.1–11.21.17

2. Argenzio E, Margadant C, Leyton-Puig D, Janssen H, Jalink K, Sonnenberg A et al. CLIC4 regulates cell adhesion and β1 integrin trafficking. J Cell Sci 2014;127:5189–203

3. Mao DY, Kleinjan ML, Jilishitz I, Swaninathan B, Obinata H, Komarova YA et al. CLIC1 and CLIC4 mediate endothelial S1P receptor signalling to facilitate Rac1 and RhoA activity and function. Sci Signal 2021;14:eabc0425

4. Peterman E, Valius M, Prekeris R. CLIC4 is a cytokinesis cleavage furrow protein that regulates cortical cytoskeleton stability during cell division. J Cell Sci 2020;133:jcs241117

5. Shukla A, Edwards R, Yang Y, Hahn A, Folkers K, Ding J et al. CLIC4 regulates TGF-β-dependent myofibroblast differentiation to produce a cancer stroma. Oncogene 2014;33:842–50

6. Ronnov-Jessen L, Villadsen R, Edwards JC, Petersen OW. Differential expression of a chloride intracellular channel gene, CLIC4, in transforming growth factor-beta1-mediated conversion of fibroblasts to myofibroblasts. Am J Pathol 2002;161:471–80

7. Wasson CW, Ross RL, Morton R, Mankouri J, Del Galdo F. The Intracellular chloride channel 4 (CLIC4) activates systemic sclerosis fibroblasts. Rheumatology (Oxford) 2020; online ahead of print

8. Shukla A, Malik M, Cataisson C, Ho Y, Friesen T, Suh KS et al. TGF-beta signalling is regulated by schnurri-2-dependent nuclear translocation of CLIC4 and consequent stabilization of phosphor-smad2 and 3. Nat Cell Biol 2009;11:777–84.

9. Yao Q, Qu X, Yang Q, Wei M, Kong B. CLIC4 mediates TGF-beta1-induced fibroblast-to-myofibroblast transdifferentiation in ovarian cancer. Oncol Rep 2009;22:541–8

10. Robbins DJ, Fei DL, Riobo NA. The hedgehog signal transduction network. Sci Signal 2012;5:re6

11. Polizio AH, Chinchilla P, Chen X, Manning DR, Riobo NA. Sonic hedgehog activates the GTPases Rac1 and RhoA in a Gli-independent manner through coupling of smoothened to Gi proteins. Sci Signal 2011;4:pt7

12. Cheng L, Owais MA, Covarrubias ML, Koch WJ, Manning DR, Peers C et al. Coupling of smoothened to inhibitory G proteins reduces voltage-gated K+ currents in cardiomyocytes and prolongs cardiac potential duration. J Biol Chem 2018;293:11022–11032

13. Yoo YA, Kang MH, Kim JS, Oh SC. Sonic hedgehog signalling promotes motility and invasiveness of gastric cancer cells through TGF-beta-mediated activation of the ALK5-Smad 3 pathway. Carcinogenesis 2008;29:480–90

14. Dennler S, Andre J, Alexaki I, Li A, Magnaldo T, ten Dijke P et al. Induction of sonic hedgehog mediators by transforming growth factor-beta: Smad3-dependent activation of Gli2 and Gli1 expression in vitro and in vivo. Cancer Res 2007;67:6981–6

15. Akhmetshina A, Palumbo K, Dees C, Bergmann C, Venalis P, Zerr P et al. Activation of canonical Wnt signalling is required for TGF-β-mediated fibrosis. Nat Commun 2012;3:735

16. Henderson Jr W, Chi EY, Ye X, Nguyen C, Tien YT, Zhou B et al. Inhibition of Wnt/beta-catenin/CREB binding protein (CBP) signalling reverses pulmonary fibrosis. Proc Natl Acad Sci U S A;107:14309–14

17. Zhou B, Liu Y, Kahn M, Ann DK, Wang H, Nguyen C et al. Interactions between β-catenin and transforming growth factor-β signaling pathways mediate epithelial-mesenchymal transition and are dependent on the transcriptional co-activator cAMP-response element-binding protein (CREB)-binding protein (CBP). J Biol Chem 2012;287:7026–38

18. Dennler S, Andre J, Verrecchia F, Mauviel A. Cloning of the human GLI2 promoter: transcriptional activation by transforming growth factor-beta via SMAD3/beta-catenin cooperation. J Biol Chem 2009;284:31523–31

19. Wasson CW, Ross RL, Wells R, Corinaldesi C, Georgiou I Ch, Riobo-Del Galdo N, Del Galdo F. Long non-coding RNA HOTAIR induces GLI2 expression through Notch signalling in systemic sclerosis dermal fibroblasts. Arthritis Res Ther;22:286

20. Gillespie J, Ross RL, Corinaldesi C, Esteves F, Derrett-Smith E, McDermott MF, et al. Transforming growth factor β activation primes canonical Wnt signalling through down-regulation of axin-2. Arthritis Rheumatol 2018;70:932–942

21. Jian H, Shen X, Liu I, Semenov M, He X, Wang XF. Smad3-dependent nuclear translocation of beta-catenin is required for TGF-beta1-induced proliferation of bone marrow-derived adult human mesenchymal stem cells. Genes Dev 2006;20:666–74.

22. DiRenzo DM, Chaudhary MA, Shi X, Franco SR, Zent J, Wang K et al. A crosstalk between TGF-β/Smad3 and Wnt/β-catenin pathways promotes vascular smooth muscle cell proliferation. Cell Signal 2016;28:498–505.

23. Walter K, Omura N, Hong SM, Griffith M, Vincent A, Borges M et al. Overexpression of smoothened activates the sonic hedgehog signalling pathway in pancreatic cancer-associated fibroblasts. Clin Cancer Res 2010;16:1781–9

24. Zhou L, Yang K, Wickett RR, Kadekaro AL, Zhang Y. Targeted deactivation of cancer-associated fibroblasts by β-catenin ablation suppresses melanoma growth. Tumour Biol 2016;37: 14235–14248

25. LeRoy EC, Medsger TA Jr. Criteria for the classification of early systemic sclerosis. J Rheumatol 2001;28:1573–6

26. Wasson CW, Abignano G, Hermes H, Malaab M, Ross RL, Jimenez SA, et al. Long non-coding RNA HOTAIR drives EZH2-dependent myofibroblast activation in systemic sclerosis through miRNA 34a-dependent activation of NOTCH. Ann Rheum Dis 2020;79:507–517

